# Distinct genetic underpinnings of inter-individual differences in the sensorimotor-association axis of cortical organisation

**DOI:** 10.1101/2023.07.13.548817

**Authors:** Giacomo Bignardi, Michel G. Nivard, H. Lina Schaare, Boris C. Bernhardt, Richard A.I. Bethlehem, Simon E. Fisher, Sofie L. Valk

## Abstract

In humans, many neurobiological features of the cortex—including gene expression patterns, microstructure, and functional connectivity—vary systematically along a sensorimotor-association (S-A) axis of brain organisation. To date, it is still poorly understood whether inter-individual differences in patterns of S-A axis capture these robust spatial relationships across neurobiological properties observed at the group-level. Here, we examine inter-individual differences in structural and functional properties of the S-A axis, namely cortical microstructure, geodesic distances, and the functional gradient, in a sample of young adults from the Human Connectome Project (N = 992, including 328 twins). We quantified heritable variation associated with inter-individual differences in the S-A axis, and assessed whether structural and functional properties that are highly spatially correlated at the group-level also share genetic underpinnings. To consider measurement errors in resting-state functional connectivity data and their impact on properties of the S-A axis, we used a multivariate twin design capable of disentangling individual-level variation in both intra- and inter-individual differences. After accounting for some of the intra-individual variation, we found average heritable individual differences in both the functional gradient (*h*_twin_^2^ = 57%), cortical microstructure (*h*_twin_^2^ = 43%), and geodesic distances (*h*_twin_^2^ = 34%). However, these genetic influences were mostly distinct and deviated from group-level patterns. In particular, we found no significant genetic correlation between the functional gradient and microstructure, while we found both positive and negative genetic associations between the functional gradient and geodesic distances. Our approach highlights the complexity of genetic contributions to brain organisation and may have potential implications for understanding cognitive variability within the S-A axis framework.

## Introduction

The human brain supports perception and action but also abstract cognition (1,2). This diversity of functions is thought to be reflected by the gradual dissociation between unimodal sensory and transmodal association cortical areas along a sensorimotor-association (S-A) axis (3). The S-A axis spans a vast array of neuroanatomical properties, including microstructural variation (myelination and cytoarchitecture) and inter-areal connectivity distance (1,3–6). Here, sensory areas show increased layer differentiation, myelination, and, predominantly, short-range connections. In contrast, association areas show less differentiated microstructural profiles, reduced myelination, and a combination of short- and long-range connectivity profiles (5,7,8).

The differentiation between sensory and association areas underwent evolutionary changes (3), with an expansion of cortical association areas paralleled by a marked laminarisation of sensory areas in human primates (9,10). Such structural re-organisation and evolutionary changes along the S-A axis (1,6,11,12) may have provided the scaffold for functional differentiation (13,14), allowing in turn for human-specific cognitive and behavioural flexibility (1,13).

Several discoveries have enhanced our understanding of links between structural and functional features of the S-A axis. These findings have highlighted spatial associations of microarchitectonic differentiation (15) and cortical geometry (16) with functional organisation (13,17). For example, T1w/T2w maps derived from non-invasive Magnetic Resonance Imaging (MRI)—indexing cortical microstructural differences —and histological markers based on cell staining have been shown to relate strongly to gene transcriptional profiles and functional dissociation along the S-A axis. Such group-level associations suggest that a canonical genetic architecture may shape S-A axis structural organisation, providing foundations for the differentiation of cortical function (6).

Recent studies noted that various features of the S-A axis show inter-individual differences in human populations that are associated with variability in a host of traits, such as neuropsychiatric traits, including autism (18), schizophrenia (19), and depression (20), as well as sex (21) and developmental differences. These findings highlight the overall importance of the S-A axis for human complex trait variation. However, it remains unclear how the different structural and functional properties of this axis relate to each other when viewed at the level of the individual, i.e. whether well-documented strong associations between such properties at the group-level (3,4,6,23–25) are also reflected in patterns of inter-individual differences. Knowing whether such patterns converge or diverge is crucial because it directly informs the study of the impact of alterations along the S-A axis on human traits. For instance, divergences between findings at the individual and group levels may suggest that associations between functional characteristics of the S-A axis and trait variability do not parallel analogous structural differences. Similarly, these differences may entail that a lack of associations between a specific neurobiological feature of the S-A axis and a particular trait does not necessarily imply the absence of relationships across other S-A axis neurobiological properties.

Here, we asked: *Do individual differences in structural properties of the S-A axis relate to differences in functional properties?* In other words, we studied whether previous widely reported group averages can inform S-A axis associations at the individual level. Specifically, we tested whether anatomical individual differences in regional cortical microstructure (6) and cortical geometry—captured by the geodesic distance of inter-connected regions across the cortical mantle (1,26)—relate to the well-known functional dissociations between sensory and transmodal association areas (1,3).

To thoroughly account for the known issue of measurement error heterogeneity across the cortex (27) and its impact on association estimates (28,29), we adapted and applied measurement error models in the form of structural equation models (30,31). This allowed us to rigorously tease apart unreliable intra-individual from reliable inter-individual variation in the functional organisation of the S-A axis.

We then went further to interrogate upstream sources of inter-individual differences in the S-A axis in structure and function: genetic variation. We asked: *Are genetic effects on the S-A axis shared across structural and functional properties?* Here, we analysed a genetically informative sample and quantified the extent of overlap across genetic effects on structural and functional properties of the S-A axis. Specifically, we used an augmented twin-informed design to quantify and tease out genetic overlaps between the S-A axis’s structural and functional properties while accounting for the impact of reliability on classic heritability estimates (31,32). Last, we evaluated the robustness of our results, both across subsamples, between regional and global cortical metrics, and between and within individuals’ S-A axis properties.

## Results

To quantify structural and functional S-A axis properties, we combined microstructural and resting-state functional MRI (rsfMRI) data from the Human Connectome Project (HCP (33); *N* = 992 adults; 529 women, mean age 28 y; 22-37 y). We computed two structural metrics and one functional metric indexing the S-A axis:

- Regional microstructure: we quantified regional microstructure indexing the differentiation between sensorimotor and association areas using the individuals’ mean intensity of regional T1w/T2w (T1w/T2w_mi_) in 400 parcels (6,34)
- Geodesic distance: we quantified cortical geometry in each individual as regional cortico-cortical network proximity (26) by computing the geodesic distance (GD) between every cortical region and its corresponding functional network, averaging within each region to get parcel-wise estimates (18). We chose geodesic distances because they align with the anatomical layout of axonal wiring along the cortical sheet and are strongly associated with functional connectivity gradients (1,35), thus providing a spatial framework that complements microstructural measures
- Functional gradient loadings: we quantified the functional S-A axis in each individual by obtaining the first component of the individual functional connectomes (FC_G1_) using diffusion map embedding (1)

We started our analysis by testing whether associations between group-level averaged maps of structural S-A axis properties correlated with the functional S-A axis (Fig. 1A-C). By using a subsample of *n* = 482 adults (229 women, mean age 28 y; 22-37 y; a subsample obtained by excluding all twins included in the full HCP sample), we were able to replicate group-level findings between averaged T1w/T2w_mi_ and FC_G1_, extending the results to GD and FC_G1_ (Fig. 1D-E).

**Figure 1.**
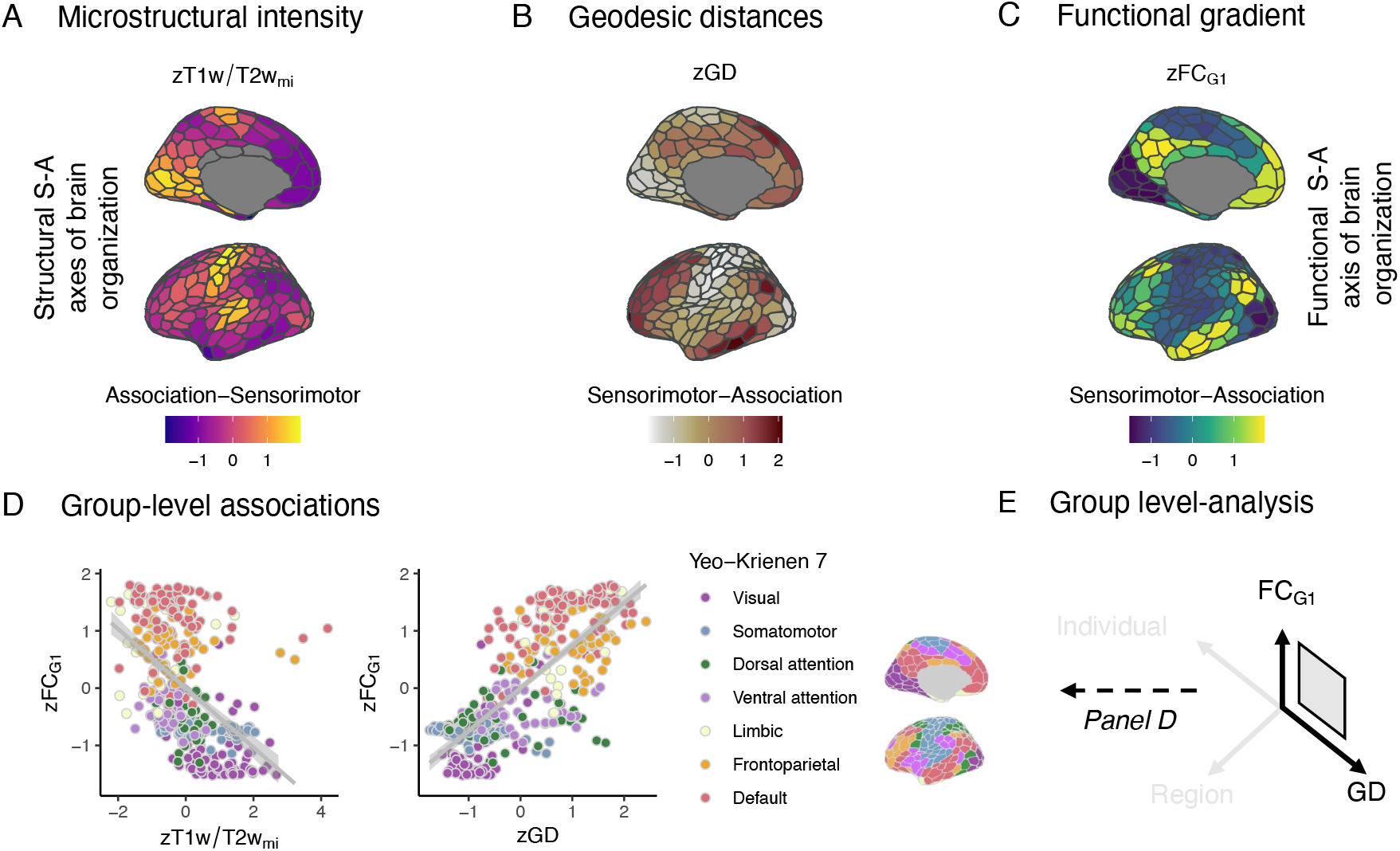
Structural and functional S-A axes strongly correlate at the group-level. Structural (**A-B**) and functional (**C**) indices of Sensorimotor-Association (S-A) axes plotted on inflated cortical surfaces (36). Values represent averages of individual T1w/T2w mean intensity profiles (A; T1w/T2w_mi_), averages of individual geodesic distances (B; GD), and functional gradients loadings (C; FC_G1_) extracted from the average of individual functional connectomes across 400 cortical regions. (**D**) Structural indices are strongly associated with functional indices of the S-A axis; Spearman 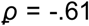 and 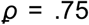 between T1w/T2w_mi_ and FC_G1_, and GD and FC_G1_, respectively; all *p* < .05. Each dot represents a regional value; the colour represents canonical Yeo-Krienen 7 network membership. (**E**) Conceptual representation of group-level analysis. Note that individual and regional information is lost in favour of group-level results.

### Pervasive inter-individual differences in the S-A axis of cortical organisation

Having estimated the extent of overlap between structural and functional S-A axis properties at the group-level, we shifted the focus to the individual level. Since group-level S-A axis can mask substantial individual variability (Fig. 2A), we asked: does individual variability in structural S-A axis properties relate to variability in S-A functional properties, as for group-level analysis (Fig. 2B)?

**Figure 2.**
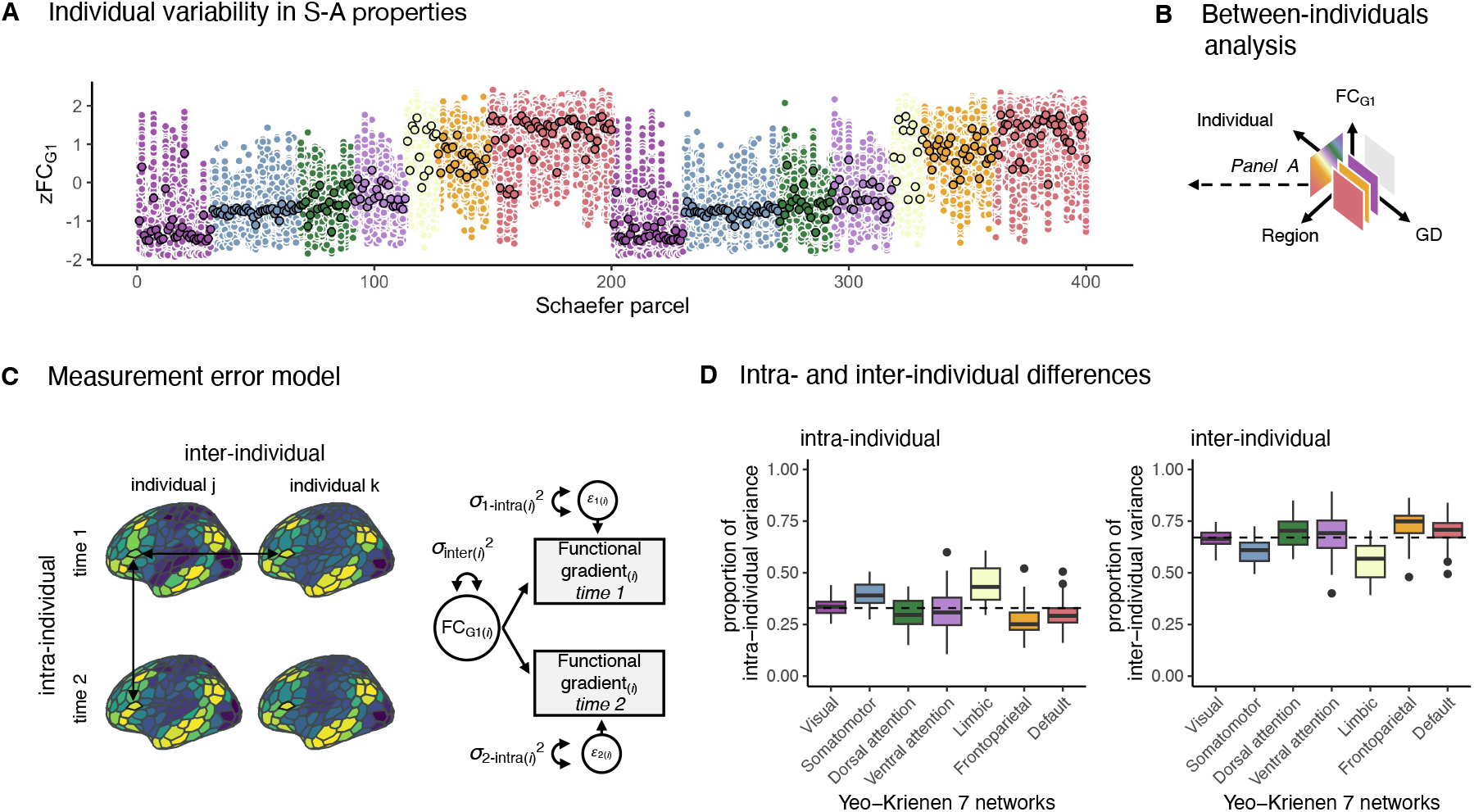
Pervasive inter-individual differences in the S-A axis of functional connectivity. (**A**) group-level estimates (black contour) overshadow pervasive individual differences in S-A axis properties. (**B**) The shift between levels of analyses: from group-level (grey square) to between-individuals (coloured squares); the gradient square conceptually captures panel A. (**C**) Measurement error model to partition, for any parcel *i*, variance in the functional gradient loadings into intra- (*σ*_d-intra(*i*)_^2^, for regional values measured at day 1 or 2 of the testing session, i.e., rectangles) and inter- (*σ*_inter(*i*)_^2^, for the latent component, i.e., circle) individual variance. Parameter estimates for any parcel *i* can be found in Supplementary Table S1. (**D**) The proportion of intra- and inter-individual variance in the functional network across Yeo-Krienen functional networks: the horizontal dashed line represents the mean proportion of variance across networks; the horizontal lines display the median within network; lower and upper hinges correspond to the first and third quartile; the whisker extends from the hinge to the largest/lower value no further than 1.5 * interquartile range from the hinge. Note that across all parcels, observed variance includes substantial inter-individual variation. *Notes on measurement model: Rectangles represent the measured phenotypes; the circle is the latent component; the double-headed arrows within the circle represent the variance associated with the latent components; one-headed arrows are the paths (here all set to 1)*.

In shifting analysis from group-level summary statistics to individual variability, we harnessed the distinction between intra-and inter-individual differences (37). The first (i.e., intra-individual) is known to index unreliable and fluctuating variability within individuals over time, while the second (i.e., inter-individual) indexes the reliable and stable part of the overall variability between individuals (Fig. 2C) (37). This distinction is crucial, as intra-individual variability can downward bias effect sizes and reduce statistical power (37), additionally downward biasing genetic estimates (38). Such factors apply heterogeneously across the whole cortex (27), and can, therefore, increase reproducibility issues (see (28) for details).

We were able to robustly distinguish between intra- and inter-individual effects by exploiting one of the strengths of the HCP design, which emphasises multiple rsfMRI sessions (across two days of scanning sessions, ∼30 min each). This feature of the HCP design allowed us to partly discard intra-individual fluctuations in rsfMRI data from inter-individual differences in the functional gradient. Precisely, we partition the inter-individual variance (*σ*inter(*i*)^2^) from the overall observed variance in the functional gradient (*σ*FCG1(*i*)^2^ for any parcel *i*) by applying a measurement error model ((30,31) Fig. 2D, see Methods).

Estimates obtained from the measurement error model indicate that 33% of the total variability in the functional gradient was, on average, accounted for by intra-individual variance (Fig. 2D) even when using individual functional gradients extracted from functional connectomes averaged across two days of rsfMRI sessions (totalling ∼60 min of scanning session). In other words, estimates for the association between the functional gradient and other S-A axis properties (or any other variable) would be, on average, biased downward by a factor of bias(*r*-observed, *r*-true) = .82 (a lower bound calculated assuming perfect reliability for the other S-A axis property (28)). Second, we observed systematic differences in estimates obtained across functional cortical networks, *F*(6, 393) = 33.21, *p* < .001; η^2^ = 0.34, 95% CI [0.27, 1.00]), with estimates for parcel-wise inter-individual variances ranging from *σ*_inter(114)_^2^= .39 to *σ*_inter(294)_^2^= .89 (Fig. 2D). That is, bias is heterogeneous and expected to influence estimates across the cortex systematically.

### Individual differences in regional cortico-cortical network proximity, rather than microstructure, relate to the functional gradient of the S-A axis of cortical organisation

To simultaneously deattenuate the heterogeneous downward biases and handle structural and functional S-A metrics, we used a Structural Equation Modelling (SEM) approach. Precisely, we specified a model in which the inter-individual differences in the functional gradient estimated via the measurement error model were directly tested for associations with microstructural profiles and geodesic distances parcel-wise data (see Methods). Here, we note that we avoided making assumptions about the causal structure generating the possible correlations between structural and functional metrics. We simply limited ourselves to estimating the association between regional properties of the S-A axis.

On the one hand, contrary to group-level topographies, we found less than 2% of the 400 parcels to display a significant association between individuals’ microstructural profiles and functional gradient loadings. These significant associations were all negative, weak (−.20 > *r* > -.27), and spread across both hemispheres and the dorsal, ventral, and default-mode functional networks. Conversely, we found large overlaps between individual geodesic distances and functional gradient loadings (Fig. 3A), with 57% of the 400 parcels showing significant associations after Bonferroni correction. The directionality of the estimates for the association between individual regional geodesic distances and functional gradient loadings highlighted systematic differences across functional networks. Significant positive associations were preferentially clustered within the visual and the default mode (one sample t-test, two-sided, *t*(21) = 5.82, *p* < .001, average *r* = .51, and *t*(53) = 4.04, *p* = .001, average *r* = .25), while negative associations were preferentially clustered within the somatomotor and ventral attention networks (one sample t-test, two-sided, *t*(59) = -13.32, *p* < .001, average *r* = -.48, and *t*(27) = -8.14, *p* < .001, average *r* = -.40, respectively, all tests accounting for multiple-testing via Bonferroni correction). Estimates obtained from standard correlation analysis further confirmed that the SEM approach successfully deattenuated measurement error bias (Fig. 3B).

**Figure 3.**
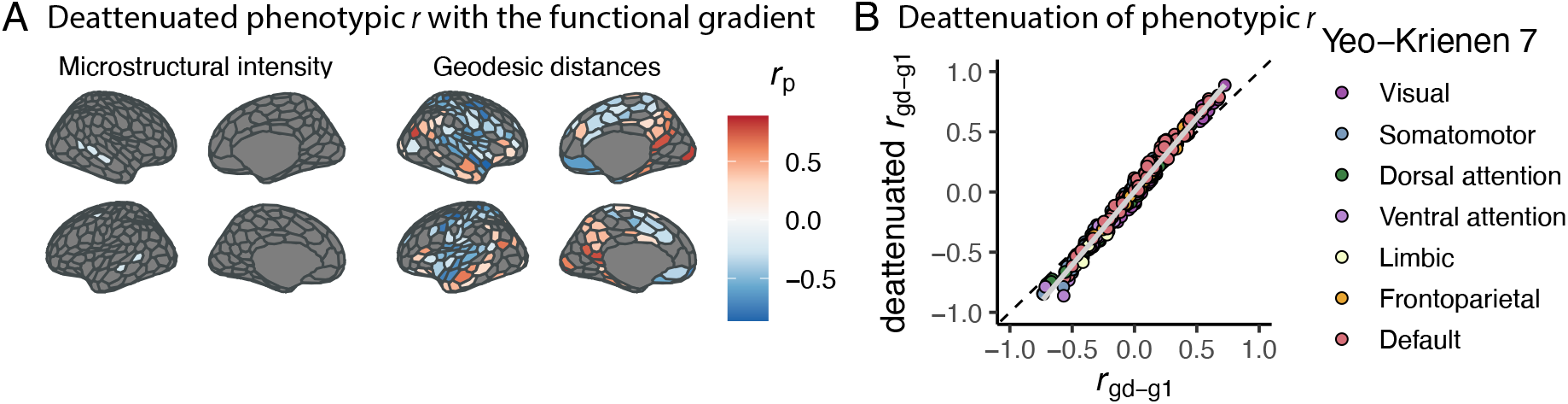
Structural and functional S-A axes selectively correlate between individuals. (**A**) Summary for the standardised estimates on the inflated cortical surface (36) from the structural measurement error model (see Methods) indicates little and weak phenotypic correlations (*r*_p_) between microstructural intensity (T1w/T2w_mi_) and functional gradient loadings (FC_G1_) inter-individual differences but large and highly significant (*p*<.05 after Bonferroni multiple-testing correction) overlaps between functional gradient loadings and geodesic distances. All parameter estimates for any parcel *i*, including the covariances between T1w/T2w_mi_ and GD that were not the focus of the current study, can be found in Supplementary Table S2. (**B**) The scatter plot shows how correlations between regional functional gradient loadings and geodesic distance estimated following the classic Pearson correlation approach (*r*_gd-g1_, on the x-axis) relate to correlations estimated with the measurement error model approach (deattenuated *r*_gd-g1_, on the y-axis). The grey line represents the deviation from the expected relationship between the two approaches under no estimated difference. As can be seen, negative and positive downward biases tend to be deattenuated.

### Genetic influences on inter-individual differences in the functional gradient of cortical organisation

We went on to assess the extent of genetic influences on the S-A axis and the degree to which their estimates are impacted by measurement error. To do so, we exploited the family structure in the HCP to partition individual differences. In particular, we applied a twin design and partitioned the variability of functional gradient loadings into genetic (*σ*_*A*_^2^; *A*: additive) and unsystematic environmental (*σ*_*E*_^2^; *E*: unique-environmental) sources via SEM. We focused our analysis on the twin HCP subsample including both monozygotic (MZ) and dizygotic (DZ) twins (*n* = 328, 195 MZ and 133 DZ individual twins, 124 and 88 women, respectively; mean age 29 y, range = 22-35 y; see Methods for details on inclusion criteria), and derived twin-based heritability estimates (*h*_twin_^2^). These *h*_twin_^2^ estimates were derived from an integrated SEM that incorporated the measurement error model. This approach made it possible to separate intra- and inter-individual differences in the functional gradient, yielding *h*_twin_^2^ estimates that robustly accounted for intra-individual variation, including measurement error.

We benchmarked our *h*_twin_^2^ estimates for functional gradient loadings (obtained from the model that accounted for measurement error) by comparing them to a classic twin model (not accounting for measurement error). Following standard cut-offs (31), we retained 395 and 398 parcel-wise measurement error and classic twin models, respectively, with a satisfactory CFI > .90 and an RMSEA < .08. We excluded an additional parcel-wise measurement error twin model as it returned an out-of-bounds estimate of *h*_twin_^2^ > 1.The inclusion of the measurement error model substantially boosted *h*_twin_^2^ estimates relative to the estimates obtained from models not accounting for intra-individual variance. In particular, accounting for intra-individual differences resulted in a much larger average heritability for functional gradient loadings: *h*_twin_^2^ = .57, *SD* = 0.13, compared to the *h*_twin_^2^ = .37, *SD* = 0.11 obtained from classic models (Fig. 4A-B). Fig. 4C provides an illustration of the enhanced estimates for the heritability of S-A axis functional organisation when properly accounting for measurement error.

**Figure 4.**
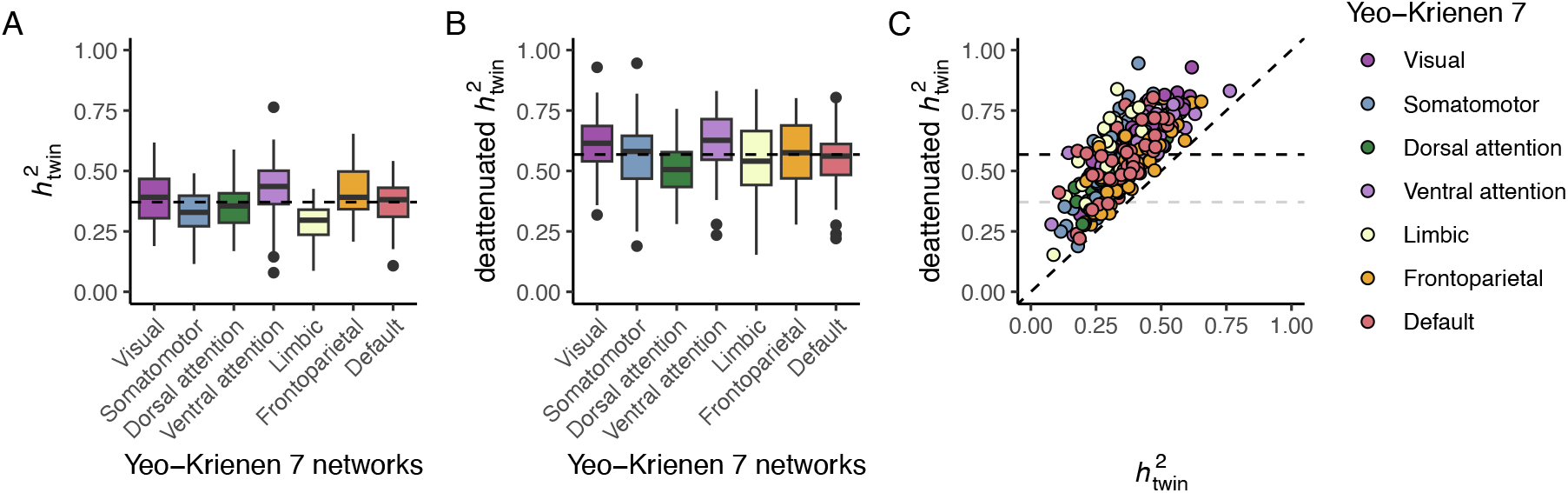
Univariate twin AE models that account for measurement error boost functional gradient heritability estimation. Parcel-wise twin heritability (*h*_twin_^2^) estimates for the 392 parcels with satisfactory fit indices across classic and measurement error twin models. **A** Box plot of the *h*_twin_^2^ stratified per Yeo-Krienen 7 functional networks. **B** Box plot of the *h*_twin_^2^ stratified per Yeo-Krienen 7 functional networks obtained from the model accounting for measurement error (here, intra-individual differences, see Methods). The dashed line displays the mean *h*_twin_^2^ across networks. Note that the average heritability is 53% higher in B. Parameter estimates for any parcel *i* can be found in Supplementary Table 3. **C** Scatter plot showing the increase of the *h*_twin_^2^ estimate across the cortex in both models. Each dot represents one parcel.

### Genetic effects on different properties of regional S-A axis variability are substantial yet mostly distinct

After having related inter-individual differences in structural and functional S-A axis properties, and showing heritable variation in functional organisation of the S-A axis, we asked whether genetic effects were mostly common or distinct across S-A axis properties. To partition variability (σ_p_ ^2^; p: S-A axis phenotypic property) within S-A modalities in *σ*_*A*_^2^ and *σ*_*E*_^2^ sources, and further unpack genetic and environmental structure-function associations, we specified a multigroup multivariate model with only A and E components (see Methods). We retained all multivariate models as they displayed satisfactory fit indices, except a model returning an out-of-bounds estimate of *h*_twin_^2^ > 1. Similar to functional gradient loadings, microstructural profiles, mean *h*_twin_^2^ = .43, *SD* = 0.11, and geodesic distances, mean *h*_twin_^2^ = .34, *SD* = 0.11, displayed substantial *h*_twin_^2^ across the cortex (Fig. 5A-B).

**Figure 5.**
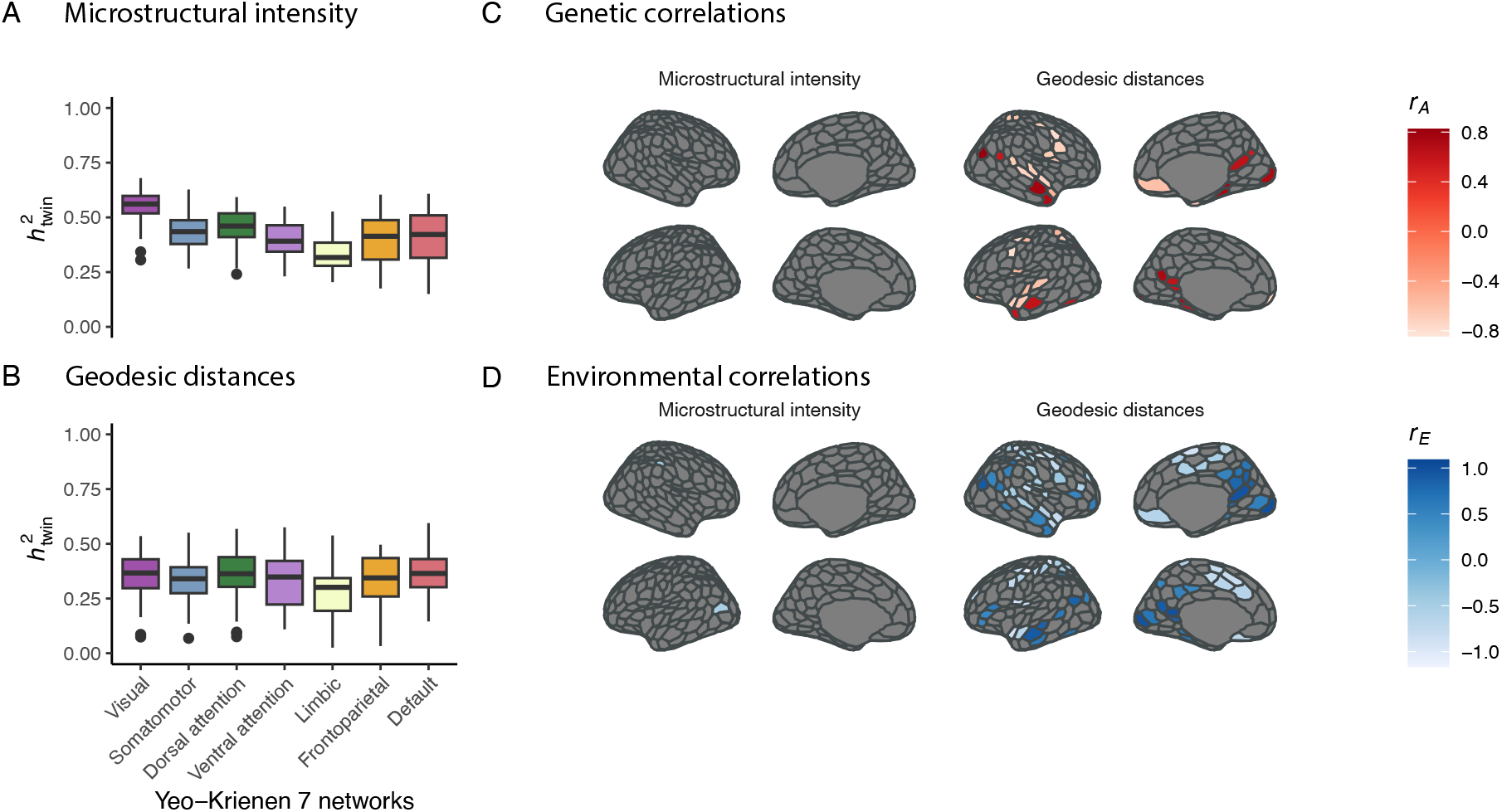
Genetic sources of structural and functional properties of the S-A axis are selectively distinct. (**A**) Parcel-wise twin heritability (*h*_twin_^2^) estimates for microstructure and (**B**) geodesic distances. (**C**) Summary for the significant additive genetic correlations *r*_A_ and (**D**) and environmental correlations (*r*_E_) across structural S-A axis properties with the functional gradient loadings on the inflated cortical surface (46). Note that the only significant genetic correlations are between geodesic distances and functional gradients loadings. The *r*_A_ estimates can be found in Supplementary Tables S4-S5.

Notwithstanding such relatively high *h*_twin_^2^ for both microstructural profiles and functional gradient loadings, we found no significant additive genetic correlation between the two (all *p* > .05, Bonferroni corrected; Fig. 5B). This suggested little room for possible shared genetic causes between the S-A axis properties indexed by microstructural intensity and functional gradient loadings. Conversely, 14% of the parcels displayed significant additive genetic correlations between geodesic distances and functional gradient loadings (7% negative and 7% positive in directionality, *p* < .05, Bonferroni corrected, Fig. 5C). The average magnitude of the genetic correlation (*r*_A_) was *r*_A_ = -.67, *SD* = 0.16, and *r*_A_ = .64, *SD* = 0.16. Furthermore, we found that for 30% of the parcels, complementary environmental effects mostly correlated between geodesic distances and functional gradient loadings (Fig. 5D).

### Associations between structural and functional properties of the S-A axis are robust across samples

To test for the robustness of the results discussed so far, we estimated the overlap of the significant regional genetic or environmental associations in the genetically informative subsample with the significant regional phenotypic associations obtained from the first subsample. Of the 104 parcels that displayed significant genetic and/or environmental correlations between geodesic distances and functional gradient loadings in the genetically informative sample, 99 also displayed a significant phenotypic correlation in the first subsample. In other words, we found a 95% overlap between subsamples in terms of the parcels implicated. These results show that regional results were robust across two subsamples drawn from the HCP.

### Genetic and environmental associations extend beyond regional S-A axis variability

As a final analysis, we asked whether associations between geodesic distances and functional gradients were generalisable beyond regional differences. First, we quantified variability in global S-A axis properties as the overall Median Absolute Deviation (MAD) across all parcels within individuals. Within an individual, higher MAD scores indicate a larger dispersion in S-A axis values across the cortex. Once more, we found no significant genetic or environmental associations between microstructural profile intensity and functional gradients MAD scores. Yet, we found a substantial negative genetic correlation between the geodesic distances and functional gradient MAD scores (*r*_*A*_ = -.78, 95% CI [-1.19, -.34], CFI = .93, RMSEA = .04; Fig. 6A).

**Figure 6.**
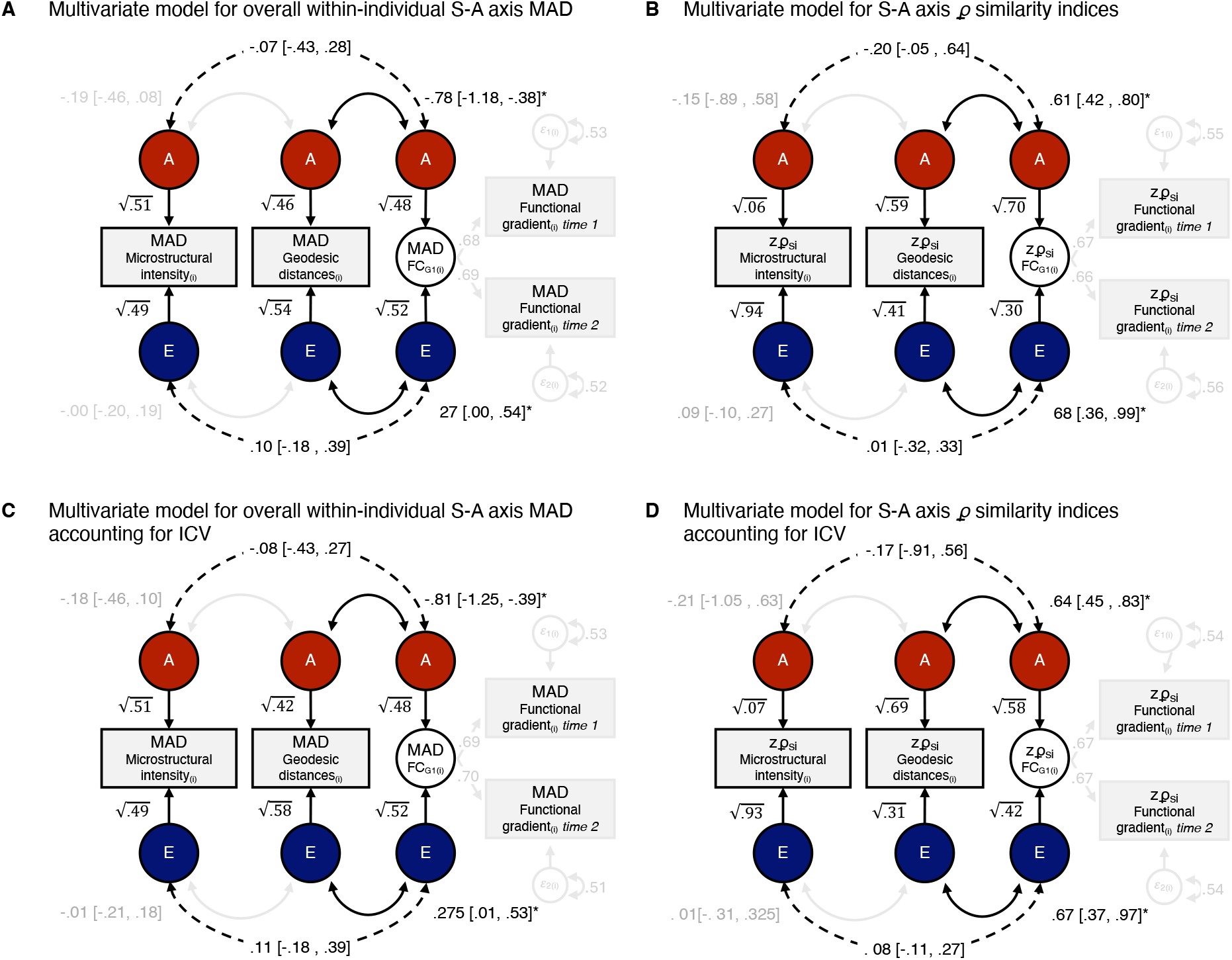
Findings extend beyond regional S-A axis associations. **(A)** Simplified graphical representation of the multivariate twin-informed SEM for overall within-individual Median Absolute Deviations (MAD). Note the strong but negative significant associations between the latent additive genetic components (*A*, circles in red) underlying geodesic distances (GD; centre) and functional gradient loadings (FC_G1_; right). The blue circles represent the latent residual environmental components (E). **(B)** Simplified graphical representation of the multivariate twin-informed SEM for 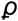 similarity indices (Fisher-*z* transformed). As for the model reported in panel A, the only significant associations are found between GD and FC_G1_. * *p* < .05. **(C)** and **(D)** panels show that the associations between MAD scores and 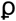 similarity indices are robust to intra-cranial volume (ICV) as a possible confounder. Here, double-headed arrows between latent variables indicate correlations (since we report standardised solutions). Dashed arrows represent nonsignificant correlations.

Additionally, to obtain a complementary estimate of global S-A axis variability, we quantified microstructural profile intensity, geodesic distance, and functional gradient 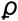 similarity indices. These indices assessed how similar S-A axis properties in one individual are compared to the average. Consistent with regional and global variance differences, 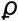 similarity indices in geodesic distance, rather than microstructure, showed strong and positive genetic correlations with global differences in the functional gradient (*r*_*A*_ = .61, 95% CI [.43, .79], CFI = .95, RMSEA = .04; Fig. 6B). Findings were robust to intra-cranial volume as a possible common cause of S-A axis structure-function covariance (Fig.6C-D)

## Discussion

Our results reveal that group-level estimates of spatial associations between structural and functional gradients differentiating sensory from association areas in the human brain might mask pervasive inter-individual differences. These inter-individual differences, in turn, might display different patterns of associations from those depicted at the group level.

Notwithstanding the comparable heritability of the different S-A axis properties as shown in this study, theoretical models of S-A axis development and evolution (3), and group-level relationships between patterns of gene expression, cortical microstructure, and functional differentiation of sensorimotor to transmodal-association areas (6), our findings revealed minimal convergence between microstructural and functional features of the S-A axis at the individual level. Specifically, we found little evidence of phenotypic and an absence of evidence for genetic associations between cortical microstructure (as measured by T1w/T2w) and S-A function (as measured by the principal gradient of functional connectivity) of the cortex.

At the same time, our results showed substantial genetic and environmental associations between individual-level differences in cortico-cortical network proximity (as measured by the geodesic distance of inter-connected regions across the cortical mantle) and S-A function. These latter results align with theories emphasising geometric constraints of brain function (1), yet do not fully align with group-level estimates. While group-level associations indicate a positive relationship between cortico-cortical network proximity, our results uncover a mixture of positive and negative relationships at the individual level of analysis (the former preferentially clustered within the visual and default mode network, the latter with the somatomotor and the ventral attention network).

Moreover, we found negative, not positive, genetic correlations when shifting from local to global association, as we did when analysing overall within-individual S-A axis dispersion. This suggests that genetic differences between people that tend to co-occur with decreased variation in geodesic distances across the cortex also tend to co-occur with more dispersed functional gradients. It is worth noting that the stronger relationship between geodesic distances and functional gradients (compared to microstructure) likely reflects the spatial constraints imposed by cortical geometry on functional organisation. While geodesic distances capture anatomical spatial relationships and functional gradients reflect connectivity patterns, both are shaped by the underlying architecture of neural connections. In particular, short-range anatomical proximity supports stronger functional coupling, which may contribute to the observed robust association. The selective associations between geodesic distances and functional gradients at the individual level also support the observed dissociation in S-A axis with microstructure, rather than reflecting a broader issue with finer spatial features of gradient-based approaches.

Fundamental principles of brain organisation can appear to be highly conserved across neurobiological properties if based on group-based analyses of topological co-variation, yet our results suggest this conservation may break down when assessed at the level of the individual. Based on group-level associations, previous work suggested that cortical maturation of diverse neurobiological properties proceeds along an evolutionary conserved and developmentally rooted S-A axis of cortical organisation (3,4). However, our results indicate that genetic variation within a population is expected to be selectively associated with some properties (e.g., function and cortico-cortical network proximity) but not others (e.g., microstructure), at least within cortical regions. The apparent paradox of observing stable group-level patterns and individual variation fits with current models of brain organisation, which recognise both shared principles and meaningful personal differences. This view is in line with recent perspectives (e.g., 39), which emphasise the importance of individual-specific brain features.

These findings may also be important for understanding the origins of differences between individuals, such as in various neuropsychiatric disorders or work on brain-behaviour associations in general. First, since we found a dissociation between structural and functional gradients, complementing studies of S-A axis variability with both structural and functional features may result in the discovery of brain-behaviour associations with selective neurobiological properties of this axis. Second, the addition of informative genetic models of structural and functional S-A axis variability would provide several novel insights into previously observed correlations. Along these lines, by disentangling brain-behaviour associations into their genetic and environmental sources, genetically informative models could also enhance our understanding of whether previously observed associations are caused by non-genetic factors. We also note that our model can be further applied to many neurobiological properties (e.g., rsfMRI, see (31)) to enhance current brain-behaviour mapping efforts (40). To facilitate this, we have made all the code available and provided all SEM functions in R and lavaan syntax.

We also underscore that the measurement error modelling approach can successfully tease apart unstable intra-individual differences from stable inter-individual differences, and this effect can have a substantial downstream impact on estimates. For example, applying the measurement error modelling approach, in line with previous results (31), resulted in a nearly 50% increase in estimates of heritability. These results should also work as a cautionary tale against interpreting differences in average heritability across principles of S-A organisation, or, more generally, between functional and structural properties of the cortex. Before accounting for intra-individual differences that occur also as a result of measurement error, heritability estimates for functional gradient loadings may have seemed lower than heritability estimates for structure. Yet, after accounting for such intra-individual variation, estimates were higher. This apparent puzzle is reconciled due to heritability being a ratio, with intra-individual variation being part of the denominator. Since functional connectivity tends to vary intra-individually, estimates will tend to be smaller when not accounting for this intra-individual variability.

We foresee that the measurement error modelling approach could have further direct application in ongoing research on the origins of psychiatric disorders and brain-behaviour studies, and in the analysis of the genomic architecture of principles of brain organisation, exactly because we show that it may mitigate the impact of measurement error heterogeneity on estimates. Indeed, when individual variability in the S-A axis is the predictor of interest, such as in brain-behaviour studies, applying any measurement error model is expected to deattenuate downwardly biased estimates (28,37,41,42). Moreover, genome-wide association studies could easily implement genome-based structural equation modelling (43,44) extensions of our approach to discard unstable and unreliable variance, overcoming attenuation biases in associations between single nucleotide polymorphisms and target phenotypes (e.g., similarly to what has been done for analyses based on polygenic indices (42)).

Yet, it is important to note that the interpretation of deattenuated estimates rests upon assumptions of what type of error is expected to influence the measurement of functional S-A axis properties across days of measurement. Here, we assumed that what is measured as being shared between sessions is the common cause of what is measured within each session. Within this framework, what is left is unique to each session, including sources of measurement error. Such application of the measurement error model to S-A axis functional properties is not expected to perfectly segregate S-A axis error-free variance in the inter-individual component (37). For example, the systematic error of individual S-A axis measurement would be indistinguishable from meaningful inter-individual variability and still be captured by the inter-individual variance component. Additionally, this measurement error model could confound unsystematic S-A axis error with genuine intra-individual differences, which may even be partially heritable (45). Therefore, the actual sources of session-to-session differences may be more nuanced than the simple measurement error model implies. Consequently, we advise caution when interpreting inter- and intra-individual differences as exclusively “error-free” and “error” variance, respectively (see Methods for further details).

Another limitation of our study is the lack of repeated structural metric measures. Although applying a measurement error model allowed us to partly disentangle intra- and inter-individual variability in functional gradient loadings, we could not account for the differences in structural properties within individuals. This limitation may have attenuated the estimated relationship between structure and function. However, the nature of the metrics and the twin design employed to elucidate differences between individuals should mitigate the impact of such a lack of repeated structural metrics (38). By applying the classical twin design, we were able to further partition unstable measurement error in the environmental (*E*) component of the model, which minimised possible biases introduced by hypothetical measurement error, at least for the additive genetic (*A*) correlations (*r*_A_) estimates.

In sum, our findings reveal that group-level results can overshadow substantial inter-individual differences within and between different neurobiological properties. By focusing on these previously underappreciated differences, we could highlight selective associations of individual variation in S-A axis cortical structure and function. These inter-individual differences and associations open a window into genetic sources of S-A axis structure and function, which we reveal to be selectively distinct. Our results underscore the complex interplay between the S-A axis’s structural and intrinsic functional properties and provide a set of tools that can be used to test their potentially differential roles in shaping cognition.

## Materials and Methods

### Sample

We used data from the Human Connectome Project (HCP) S1200 release. The HCP includes data from 1206 individuals (656 women) that comprise 298 Monozygotic (MZ) twins, 188 Dizygotic (DZ) twins, and 720 individuals, with mean age ± *SD* = 28.8 ± 3.7 years (age range = 22-37 years). Informed consent for all individuals was obtained by HCP, and our data usage was approved by HCP and complied with all relevant ethical regulations for working with human participants (see (13,33,46)). The primary participant pool comes from individuals born in Missouri to families that include twins, sampled as healthy representatives of ethnic and socioeconomic diversity of US individuals, based on data from the Missouri Department of Health and Senior Services Bureau of Vital Records. We followed standard guidelines for inclusion criteria as described elsewhere (13). Our sample, in line with Valk et al., (13) comprised 992 (529 women) individuals. The first subsample of *n* = 482 (229 women) was created by excluding all twins. The second genetically informative subsample of *n* = 328 (212 women) was created by including only twins with genotyped zygosity matching self-reported zygosity (195 MZ and 133 DZ individuals; 124 women and 88 men, respectively, forming between 150 and 152 complete pairs, depending on data availability for the S-A axis modality).

### Functional imaging

Functional connectivity matrices were based on four 14 min 33 s of functional Magnetic Resonance Imaging (fMRI) data acquired over two sessions, spaced two days apart, through the HCP, which underwent HCP’s minimal preprocessing. No global signal regression was performed in the processing of the fMRI data. For each individual, four functional connectivity matrices were computed using the minimally preprocessed, spatially normalised resting-state fMRI (rsfMRI) scans, which were co-registered using MSMAll to template HCP 32k_LR surface space. 32k_LR surface space consists of 32,492 total nodes per hemisphere (59,412 excluding the medial wall). We computed four functional connectivity matrices per individual from the average time series extracted in each of the 400 Schaefer cortical parcels. The individual functional connectomes were generated by averaging preprocessed time series within nodes, Pearson correlating nodal time series and converting them to Fisher-*z* scores. The average functional connectomes were obtained by averaging functional connectomes within individuals (i.e., between sessions) and between individuals.

### Structural imaging

MRI protocols of the HCP have been previously described (33,46). MRI data were acquired originally on the same day on the HCP’s custom 3T Siemens Skyra equipped with a 32-channel head coil. T1w images with identical parameters were acquired using a 3D-MP-RAGE sequence over 7 min 40 s (0.7 mm isovoxels, matrix = 320 × 320, 256 sagittal slices; TR = 2400 ms, TE = 2.14 ms, TI = 1000 ms, flip angle = 8°; iPAT = 2). T2w images were acquired using a 3D T2-SPACE sequence with identical geometry over 8 min and 24 s (TR = 3200 ms, TE = 565 ms, variable flip angle; iPAT = 2). We followed the preprocessing steps outlined in Valk et al. (13).

### Parcellation and functional networks

We used the Schaefer group-level hard-parcellation, originally obtained by a gradient-weighted Markov random field model integrating local gradient and global similarity approaches (34). To stratify results within canonical cortical functionally coupled networks, we used the seven Yeo-Krienen networks (47).

### Microstructural profiles (T1w/T2w_mi_)

We used T1w/T2w imaging myelin-sensitive contrast from the HCP minimal processing pipeline, which uses the T2w to correct for inhomogeneities in the T1w image to estimate mean intensity T1w/T2w microstructural profiles (T1w/T2w_mi_). T1w/T2w_mi_ has been shown to map to model-based tract-tracing histological data in macaque, estimate intracortical myelin content, and thus approximate architectural complexity and cortical hierarchy (6).

### Geodesic distance (GD)

Individual geodesic distances (GD) were computed using the Micapipe toolbox (26). Briefly, we computed GD between each region and their top 10% of maximally functionally connected regions along each individual native cortical midsurface. We further averaged within regions to obtain a parcel-wise value and improve computation performance. Micapipe implements the Dijkstra algorithm (48) (further details can be found in (26)).

### Functional gradient loadings (FC_G1_)

We sequentially averaged FCs, first within days, resulting in two FCs per individual, and then between days, resulting in one FC per individual. We then extracted the three first components from the two sequentially averaged and one averaged FCs, using the Python package BrainSpace (49). Extraction of the first eigenvector followed standard procedures, with the original individual FCs set at a connection density of 10% (i.e., the FCs were made sparse by setting a sparsity threshold of 90%). The first ten eigenvectors were then obtained by decomposing the FCs by diffusion map embedding, a robust non-linear manifold learning technique (1). To aid comparability across individuals, we aligned individual eigenvectors to the template eigenvector by Procrustes rotation (50). The template functional gradient was directly extracted from the overall mean FC matrix.

### Group-level associations analysis

We computed Spearman rank-order correlations 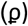 between the structural (T1w/T2w_mi_ and GD) and functional (FC_G1_) S-A axis group-level properties. Group-level properties were obtained from the average of the individual structural S-A properties (i.e., average T1w/T2w_mi_ and GD), and from the decomposition of the average FC (i.e., principal gradient obtained via diffusion map embedding of the average FC).

### Measurement model of error in individual variability of the functional S-A axis

To partition stable inter-individual variability in functional gradient loading, we adapted previous measurement error models to rsfMRI to the functional gradient (30,31). The intuition behind such a modelling strategy is simple. Suppose parcel-wise values are measured without error and are stable over a reasonable period of time (e.g., one day). In that case, the correlations across individuals between the values obtained across two time points will equal 1. If the correlations deviate from 1 instead, regional values will be measured with some error, with bigger deviations corresponding to higher error or fluctuation over time. As can be seen in Fig. 2C, when errors or changes over time are present, it may become difficult to distinguish regional differences between individuals from regional differences within individuals. In this case, we can use the measurement error model to estimate what stays constant across time, indexing the “true” regional values. Across the manuscript, for correctness, since “error” variance can include meaningful, yet unstable, fluctuation in rsfMRI, while “true” variance can also consist of systematic measurement error across sessions, we refer to the former term as intra-individual and the latter as inter-individual variability (37). First, we fit a measurement model to parcel-wise functional gradient loadings averaged within days. In line with Teeuw et al. (31), we did not constrain intra-individual variance components to be equal across days of scanning sessions. We performed model fitting in lavaan (44) after standardising observed variables (i.e., std.ov = T). We then used model estimates obtained for the variances of the latent and observed components. Using Spearman-Brown correction, we computed the averaged proportion of stable inter-individual variance in functional gradient loadings across days as the intra-class correlation (ICC) (51). For each parcel *i*, the ICC was calculated as follows:

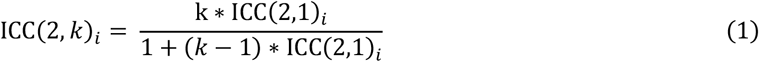

Where *k* is a constant equal to the number of measures (i.e., *k* = 2) and the ICC(2,1)_*i*_ is calculated as follows:

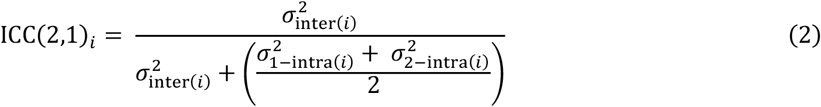

ICC(2,*k*)_*i*_ estimates the proportion of inter-individual variance over the total variance, *σ*_inter(*i*)_^2^, in the functional loadings as if they were obtained from the average of the two scanning sessions. The proportion of intra-individual variance for a parcel *i, σ*_intra(*i*)_^2^, is obtained simply by subtracting the ICC(2,*k*)_*i*_ from 1.

The expected bias for any parcel *i* was calculated following Tiego et al. (28):

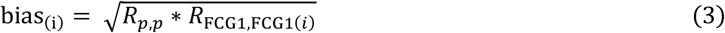

Where *R*_*p,p*_, the reliability for the structural S-A axis property *p* (e.g., T1w/T2w_mi_), was set to be equal to 1 across all parcels, and *R*_FCG1,FCG1(*i*)_, the reliability of parcel-wise value for the functional gradient loading, was calculated as ICC(2,*k*)_*i*_.

### Multivariate Measurement error SEM

We used Structural Equation Modelling (SEM) to estimate correlations between structural (i.e., T1w/T2w_mi(*i*)_, GD_(*i*)_) and functional (i.e, FC_G1(*i*)_) S-A modalities. Each multivariate model simultaneously accounted for intra-individual variances by including the measurement error model (Fig. 7A). All models were fitted in lavaan (52) after standardising all observed variables (i.e., std.ov = T). Prior to model fitting, sex and age were regressed from parcel-wise S-A axis values using the function umx::umx_residualize() (53). Structural equation models were fit to residual scores. We assumed missing data to be missed at random and followed parameters’ estimation via full-information Maximum Likelihood (i.e., missing = “ML”).

**Figure 7.**
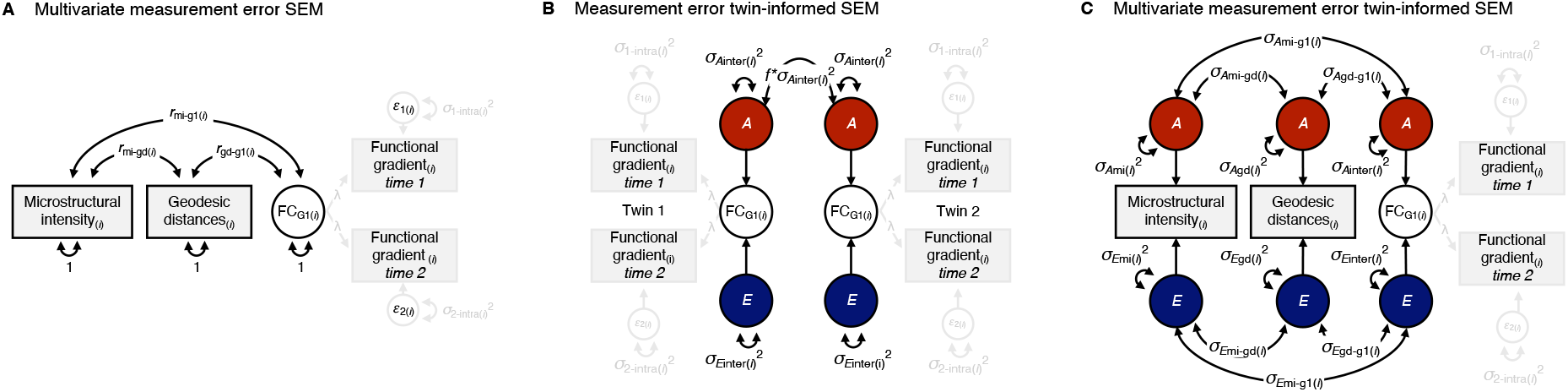
SEM approach. (**A**) Structural measurement error model used to estimate deattenuated correlation between S-A axis properties considering the distinction between intra- and inter-individual differences in functional gradient loading variability. Observed parcel-wise values and latent components are standardised before model fitting. (**B**) Twin model used to obtain deattenuated *h*_twin_^2^ estimates. Here, the covariance between the twins’ parcel-wise functional gradient loadings is estimated directly on the inter-individual component of variance (i.e., FC_G1_) and set to *f* * *σ*_*A*inter_^2^, where *f* = 1 for MZ twins and f = ½ for DZ twins. The model assumes absence of shared household and non-additive genetic effects on functional gradients in young adults. (**C**) Simplified graphical representation of the multivariate twin-informed SEM, which puts the models depicted in A and B together. *Notes on SEM: Rectangles represent the measured structural or functional MRI-derived phenotypes; the circle is the latent components; the double-headed arrows connecting circles with themselves represent the variance associated with the latent components; double-headed arrows connected between circles covariances; Where not noted, one-headed arrows are the paths (here all set to 1)*.

### Twin-informed Multivariate Structural Equation Modelling

We used multigroup SEM to partition parcel-wise variability in structural (*σ*T1w/T2w_mi(*i*)_^2^, *σ*GD_(*i*)_^2^) and functional (*σ*_inter(*i*)_^2^) S-A modalities into either additive genetic (*σ*_*A*_^2^) or unsystematic environmental (*σ*_*E*_^2^) sources of variance. Structural equation models were fit to T1w/T2w_mi*i*_, GD_*i*_, and FC_G1*i*1_ (day 1) and FC_G1*i*2_ (day 2) data, grouped by zygosity (i.e., two groups). The model specification was informed by the multivariate twin design (54). Briefly, monozygotic (MZ) twins are ∼100 % genetically identical, coming from the same fertilised egg. In contrast, dizygotic (DZ) twins are, on average, only 50% additively genetically similar regarding allelic variants coming from two different fertilised eggs. Thus, the correlation *f* between the additive genetic component (A) is set to be equal to 1 for MZ and ½ for DZ. In contrast, each twin’s unique environment (E) component will be unique; therefore, their correlation will be equal to 0. In total, we fit one multivariate AE model per parcel. Following the measurement error procedure outlined above and in reference (31), a common pathway measurement error model was included in the specification of the multigroup SEM (Fig. 7B). As such, each multivariate model simultaneously accounted for intra-individual variance (Fig. 7C). We employed the direct symmetric approach by estimating variance components directly while setting path coefficients to 1 (with the exception of the measurement model, for which we fixed the variance to be equal to 1, and estimated the path coefficients, instead). We chose this approach as it has been shown to reduce type I errors and produce asymptotically unbiased χ^2^ (55).

Similarly to reference (56), twin models were fitted in lavaan (52), with standardisation of observed variables before model fitting (i.e., std.ov = T). To control for the effect of age and sex on S-A axis properties, we residualised parcel-wise variables prior to modelling using the function umx::umx_residualize() (53). Residuals were used as observed variables in later twin modelling. We estimated parameters via full-information Maximum Likelihood (i.e., missing = “ML”) and evaluated the goodness of fit for each parcel by comparative fit index (CFI) and root mean square error of approximation (RMSEA) scores. Following standard cut-offs (31), we retained only models with a “satisfactory” CFI>.90 and an RMSEA<.08. Narrow-sense twin heritability (*h*_twin_^2^) estimates for each parcel *i* were defined as the ratio of the additive genetic variance over the sum of the additive genetic and environmental variances:

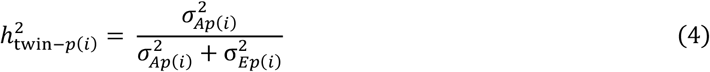

Where *σ*_*Ap(i)*_ ^2^ is the additive genetic variance for the S-A axis parcel-wise value for the given property *p* (e.g., GD). For functional gradient loadings, the heritability was calculated as

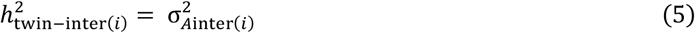

After imposing the equality constraint on the common factor FC_G1*inter(i)*_

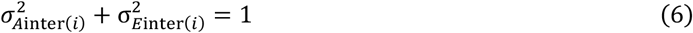

Genetic correlations (*r*_A_) were calculated as:

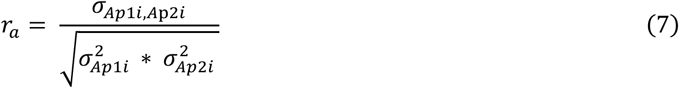

Where *σ*_*Ap*1,*Ap*2_^2^ is the additive genetic covariance between two S-A axis properties, *p*1 and *p*2 (e.g., GD and T1w/T2w_mi_). Environmental correlations (*r*_*E*_) were calculated similarly to *r*_*A*_ but using environmental variance and covariance estimates.

### Generalisation beyond regional associations

For each individual, we obtain two metrics for structural and functional S-A axis properties (i.e., a total of six measures per individual):

*Overall within-individual Median Absolute Deviation*: we quantified the spread of the regional values across the cortex by computing within-individual Median Absolute Deviation (MAD) of microstructure, geodesic distances, and functional gradient loadings. MAD is a robust univariate measure of statistical dispersion and is simply calculated as follows:

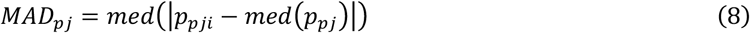

Where *p*_*ij*_ is the parcel-wise value for a property *p*, an individual *j*, and a parcel *I* and *med* is the median.

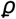 *similarity index*: we obtained the similarity index by estimating the Spearman rank (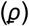) correlations between each individual microstructure, cortico-cortical network proximity, and functional gradient loadings with the respective S-A group-level modality vectors. For example, the 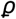 similarity index for the cortico-cortical network proximity for an individual *j* was obtained by correlating their GD with the group-level GD. Similarly, for the same individual *j*, the similarity index for their functional gradient loading was obtained by correlating their FC_G1_ on day 1 and on day 2 of scanning with the group-level FC_G1_.

Similar to what is outlined above for regional analysis, we fit two multivariate AE models, one per metric. Before model fitting, 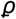 similarity indices were first Fisher-*z* transformed. To recapitulate regional analysis as closely as possible within the multivariate model, we also included the measurement error model to overall within-individual MAD and 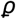 similarity index functional gradient loadings obtained on days 1 and 2 of scanning sessions. Note that standardised coefficients are obtained using the lavaan::standardizedSolution() function. As a final sensitivity analysis, to discount individuals’ whole brain volume as a possible confounding effect of the relationship between SA axis structure-function associations, we additionally included total intra-cranial volume. Precisely, we followed a two-step procedure to discount intra-cranial volume as a possible common cause. First, we regressed out intra-cranial volume from overall within-individual MAD and Fisher-*z* transformed 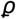 similarity indices for all S-A axis properties. We then re-fit the exact multivariate twin models to the residuals.

## Data availability

We obtained human data from the open-access Human Connectome Project (HCP) S1200 young adult sample. HCP Young Adult data are available at https://www.humanconnectome.org/study/hcp-young-adult. The Supplementary Table with summary statistics can be found at https://github.com/giacomobignardi/h2_SA_axis/tree/main/SI.

## Code availability

All code is available and can be found at https://github.com/giacomobignardi/h2_SA_axis. SEM and twin-based analysis were carried out using the statistical package latent variable analysis (lavaan) https://lavaan.ugent.be/. The function to apply the measurement error model (meermo) can be found here: https://github.com/giacomobignardi/h2_SA_axis/tree/main/R/functions/meermo. The lavaan syntax for latent variable analysis of twin data (lavaantwda) can be found in the repository https://github.com/giacomobignardi/h2_SA_axis/tree/main/R/functions/lavaantwda. An introduction to twin modelling using lavaan can be found at https://rpubs.com/MichelNivard/798608. Code and tutorial for functional gradient decomposition of functional connectomes are available at https://brainspace.readthedocs.io/en/latest/pages/install.html. The code and tutorial to generate geodesic distances can be found at https://micapipe.readthedocs.io/en/latest/.

## Acknowledgements

We want to thank the Human Connectome Project, Washington University, the University of Minnesota, and Oxford University Consortium (Principal Investigators: David Van Essen and Kamil Ugurbil; 1U54MH091657) originally funded by the 16 N.I.H. Institutes and Centers that support the N.I.H. Blueprint for Neuroscience Research; and by the McDonnell Center for Systems Neuroscience at Washington University. We would also like to thank Jitse Amelink, Meike D. Hettwer, and MacKenzie D. Trupp for their comments on an earlier version of the manuscript. This study was supported by the German Federal Ministry of Education and Research (BMBF), the Max Planck Society, and the Max Planck School of Cognition.

## References

1. Margulies DS, Ghosh SS, Goulas A, Falkiewicz M, Huntenburg JM, Langs G, et al. Situating the default-mode network along a principal gradient of macroscale cortical organization. Proc Natl Acad Sci. 2016 Nov 1;113(44):12574–9.

2. Mesulam MM. From sensation to cognition. Brain. 1998 Jun 1;121(6):1013–52.

3. Sydnor VJ, Larsen B, Bassett DS, Alexander-Bloch A, Fair DA, Liston C, et al. Neurodevelopment of the association cortices: Patterns, mechanisms, and implications for psychopathology. Neuron. 2021 Sep 15;109(18):2820–46.

4. Sydnor VJ, Larsen B, Seidlitz J, Adebimpe A, Alexander-Bloch AF, Bassett DS, et al. Intrinsic activity development unfolds along a sensorimotor–association cortical axis in youth. Nat Neurosci. 2023 Apr;26(4):638–49.

5. von Economo CF, Koskinas GN. Die cytoarchitektonik der hirnrinde des erwachsenen menschen. J Springer. 1925;

6. Burt JB, Demirtaş M, Eckner WJ, Navejar NM, Ji JL, Martin WJ, et al. Hierarchy of transcriptomic specialization across human cortex captured by structural neuroimaging topography. Nat Neurosci. 2018 Sep;21(9):1251–9.

7. Brodmann K. ergleichende Lokalisationslehre der Grosshirnrinde in ihren Prinzipien dargestellt auf Grund des Zellenbaues. Barth. 1909;

8. Ercsey-Ravasz M, Markov NT, Lamy C, Van Essen DC, Knoblauch K, Toroczkai Z, et al. A predictive network model of cerebral cortical connectivity based on a distance rule. Neuron. 2013 Oct 2;80(1):184–97.

9. Buckner RL, Krienen FM. The evolution of distributed association networks in the human brain. Trends Cogn Sci. 2013 Dec 1;17(12):648–65.

10. García-Cabezas MÁ, Hacker JL, Zikopoulos B. Homology of neocortical areas in rats and primates based on cortical type analysis: an update of the Hypothesis on the Dual Origin of the Neocortex. Brain Struct Funct. 2023 Jun 1;228(5):1069–93.

11. Saberi A, Paquola C, Wagstyl K, Hettwer MD, Bernhardt BC, Eickhoff SB, et al. The regional variation of laminar thickness in the human isocortex is related to cortical hierarchy and interregional connectivity. PLoS Biol. 2023 Nov;21(11):e3002365.

12. Markov NT, Ercsey-Ravasz M, Van Essen DC, Knoblauch K, Toroczkai Z, Kennedy H. Cortical high-density counterstream architectures. Science. 2013 Nov 1;342(6158):1238406.

13. Valk SL, Xu T, Paquola C, Park B yong, Bethlehem RAI, Vos de Wael R, et al. Genetic and phylogenetic uncoupling of structure and function in human transmodal cortex. Nat Commun. 2022 May 9;13(1):2341.

14. Demirtaş M, Burt JB, Helmer M, Ji JL, Adkinson BD, Glasser MF, et al. Hierarchical Heterogeneity across Human Cortex Shapes Large-Scale Neural Dynamics. Neuron. 2019 Mar 20;101(6):1181-1194.e13.

15. Paquola C, Amunts K, Evans A, Smallwood J, Bernhardt B. Closing the mechanistic gap: the value of microarchitecture in understanding cognitive networks. Trends Cogn Sci. 2022 Oct 1;26(10):873–86.

16. Leech R, Vos De Wael R, Váša F, Xu T, Austin Benn R, Scholz R, et al. Variation in spatial dependencies across the cortical mantle discriminates the functional behaviour of primary and association cortex. Nat Commun. 2023 Sep 13;14(1):5656.

17. Suárez LE, Markello RD, Betzel RF, Misic B. Linking Structure and Function in Macroscale Brain Networks. Trends Cogn Sci. 2020 Apr 1;24(4):302–15.

18. Hong SJ, Vos de Wael R, Bethlehem RAI, Lariviere S, Paquola C, Valk SL, et al. Atypical functional connectome hierarchy in autism. Nat Commun. 2019 Mar 4;10(1):1022.

19. Dong D, Yao D, Wang Y, Hong SJ, Genon S, Xin F, et al. Compressed sensorimotor-to-transmodal hierarchical organization in schizophrenia. Psychol Med. 2023 Feb;53(3):771–84.

20. Xia M, Liu J, Mechelli A, Sun X, Ma Q, Wang X, et al. Connectome gradient dysfunction in major depression and its association with gene expression profiles and treatment outcomes. Mol Psychiatry. 2022 Mar;27(3):1384–93.

21. Serio B, Hettwer MD, Wiersch L, Bignardi G, Sacher J, Weis S, et al. Sex differences in functional cortical organization reflect differences in network topology rather than cortical morphometry. Nat Commun. 2024 Sep 4;15(1):7714.

22. Dong HM, Margulies DS, Zuo XN, Holmes AJ. Shifting gradients of macroscale cortical organization mark the transition from childhood to adolescence. Proc Natl Acad Sci. 2021 Jul 13;118(28):e2024448118.

23. Huntenburg JM, Bazin PL, Margulies DS. Large-Scale Gradients in Human Cortical Organization. Trends Cogn Sci. 2018 Jan 1;22(1):21–31.

24. Zhang XH, Anderson KM, Dong HM, Chopra S, Dhamala E, Emani PS, et al. The cell-type underpinnings of the human functional cortical connectome. Nat Neurosci. 2024 Nov 21;1–11.

25. Goulas A, Changeux JP, Wagstyl K, Amunts K, Palomero-Gallagher N, Hilgetag CC. The natural axis of transmitter receptor distribution in the human cerebral cortex. Proc Natl Acad Sci. 2021 Jan 19;118(3):e2020574118.

26. Cruces RR, Royer J, Herholz P, Larivière S, Vos de Wael R, Paquola C, et al. Micapipe: A pipeline for multimodal neuroimaging and connectome analysis. NeuroImage. 2022 Nov 1;263:119612.

27. Helwegen K, Libedinsky I, Heuvel MP van den. Statistical power in network neuroscience. Trends Cogn Sci. 2023 Mar 1;27(3):282–301.

28. Tiego J, Martin EA, DeYoung CG, Hagan K, Cooper SE, Pasion R, et al. Precision behavioral phenotyping as a strategy for uncovering the biological correlates of psychopathology. Nat Ment Health. 2023 May;1(5):304–15.

29. Kang K, Seidlitz J, Bethlehem RAI, Xiong J, Jones MT, Mehta K, et al. Study design features increase replicability in brain-wide association studies. Nature. 2024 Nov 27;1–9.

30. Brandmaier AM, Wenger E, Bodammer NC, Kühn S, Raz N, Lindenberger U. Assessing reliability in neuroimaging research through intra-class effect decomposition (ICED). Johansen-Berg H, Kastner S, editors. eLife. 2018 Jul 2;7:e35718.

31. Teeuw J, Hulshoff Pol HE, Boomsma DI, Brouwer RM. Reliability modelling of resting-state functional connectivity. NeuroImage. 2021 May 1;231:117842.

32. Ge T, Holmes AJ, Buckner RL, Smoller JW, Sabuncu MR. Heritability analysis with repeat measurements and its application to resting-state functional connectivity. Proc Natl Acad Sci. 2017 May 23;114(21):5521–6.

33. Van Essen DC, Ugurbil K, Auerbach E, Barch D, Behrens TEJ, Bucholz R, et al. The Human Connectome Project: A data acquisition perspective. NeuroImage. 2012 Oct;62(4):2222–31.

34. Schaefer A, Kong R, Gordon EM, Laumann TO, Zuo XN, Holmes AJ, et al. Local-Global Parcellation of the Human Cerebral Cortex from Intrinsic Functional Connectivity MRI. Cereb Cortex. 2018 Sep 1;28(9):3095–114.

35. Oligschläger S, Huntenburg JM, Golchert J, Lauckner ME, Bonnen T, Margulies DS. Gradients of connectivity distance are anchored in primary cortex. Brain Struct Funct. 2017 Jul;222(5):2173– 82.

36. Mowinckel AM, Vidal-Piñeiro D. Visualization of Brain Statistics With R Packages ggseg and ggseg3d. Adv Methods Pract Psychol Sci. 2020 Dec 1;3(4):466–83.

37. Zuo XN, Xu T, Milham MP. Harnessing reliability for neuroscience research. Nat Hum Behav. 2019 Aug;3(8):768–71.

38. Hagenbeek FA, Hirzinger JS, Breunig S, Bruins S, Kuznetsov DV, Schut K, et al. Maximizing the value of twin studies in health and behaviour. Nat Hum Behav. 2023 Jun;7(6):849–60.

39. Petersen SE, Seitzman BA, Nelson SM, Wig GS, Gordon EM. Principles of cortical areas and their implications for neuroimaging. Neuron. 2024 Sep 4;112(17):2837–53.

40. Marek S, Tervo-Clemmens B, Calabro FJ, Montez DF, Kay BP, Hatoum AS, et al. Reproducible brain-wide association studies require thousands of individuals. Nature. 2022 Mar;603(7902):654–60.

41. Xu T, Kiar G, Cho JW, Bridgeford EW, Nikolaidis A, Vogelstein JT, et al. ReX: an integrative tool for quantifying and optimizing measurement reliability for the study of individual differences. Nat Methods. 2023 Jul;20(7):1025–8.

42. van Kippersluis H, Biroli P, Dias Pereira R, Galama TJ, von Hinke S, Meddens SFW, et al. Overcoming attenuation bias in regressions using polygenic indices. Nat Commun. 2023 Jul 25;14(1):4473.

43. Pritikin JN, Neale MC, Prom-Wormley EC, Clark SL, Verhulst B. GW-SEM 2.0: Efficient, Flexible, and Accessible Multivariate GWAS. Behav Genet. 2021 May;51(3):343–57.

44. Grotzinger AD, Rhemtulla M, de Vlaming R, Ritchie SJ, Mallard TT, Hill WD, et al. Genomic structural equation modelling provides insights into the multivariate genetic architecture of complex traits. Nat Hum Behav. 2019 May;3(5):513–25.

45. Kemper KE, Sidorenko J, Wang H, Hayes BJ, Wray NR, Yengo L, et al. Genetic influence on within-person longitudinal change in anthropometric traits in the UK Biobank. Nat Commun. 2024 May 6;15(1):3776.

46. Glasser MF, Sotiropoulos SN, Wilson JA, Coalson TS, Fischl B, Andersson JL, et al. The minimal preprocessing pipelines for the Human Connectome Project. NeuroImage. 2013 Oct 15;80:105– 24.

47. Yeo BTT, Krienen FM, Sepulcre J, Sabuncu MR, Lashkari D, Hollinshead M, et al. The organization of the human cerebral cortex estimated by intrinsic functional connectivity. J Neurophysiol. 2011 Sep;106(3):1125–65.

48. Dijkstra EW. A note on two problems in connexion with graphs. Numer Math. 1959 Dec 1;1(1):269–71.

49. Vos de Wael R, Benkarim O, Paquola C, Lariviere S, Royer J, Tavakol S, et al. BrainSpace: a toolbox for the analysis of macroscale gradients in neuroimaging and connectomics datasets. Commun Biol. 2020 Mar 5;3(1):1–10.

50. Bethlehem RAI, Paquola C, Seidlitz J, Ronan L, Bernhardt B, Consortium CC, et al. Dispersion of functional gradients across the adult lifespan. NeuroImage. 2020 Nov 15;222:117299.

51. Koo TK, Li MY. A Guideline of Selecting and Reporting Intraclass Correlation Coefficients for Reliability Research. J Chiropr Med. 2016 Jun;15(2):155–63.

52. Rosseel Y. lavaan: An R Package for Structural Equation Modeling. J Stat Softw. 2012 May 24;48(1):1–36.

53. Bates TC, Maes H, Neale MC. umx: Twin and Path-Based Structural Equation Modeling in R. Twin Res Hum Genet Off J Int Soc Twin Stud. 2019 Feb;22(1):27–41.

54. Boomsma D, Busjahn A, Peltonen L. Classical twin studies and beyond. Nat Rev Genet. 2002 Nov;3(11):872–82.

55. Verhulst B, Prom-Wormley E, Keller M, Medland S, Neale MC. Type I Error Rates and Parameter Bias in Multivariate Behavioral Genetic Models. Behav Genet. 2019 Jan;49(1):99–111.

56. Bignardi G, Wesseldijk LW, Mas-Herrero E, Zatorre RJ, Ullén F, Fisher SE, et al. Twin modelling reveals partly distinct genetic pathways to music enjoyment. Nat Commun. 2025 Mar 25;16(1):2904.

